# Conserved role of spike S2 domain N-glycosylation across beta-coronavirus family

**DOI:** 10.1101/2024.09.05.611372

**Authors:** Qi Yang, Anju Kelkar, Balaji Manicassamy, Sriram Neelamegham

**Affiliations:** Chemical & Biological Engineering, State University of New York, Buffalo, NY 14260, USA; Cell, Gene and Tissue Engineering Center, State University of New York, Buffalo, NY 14260, USA; Microbiology and Immunology, University of Iowa, Iowa City, IA 52242, USA; Biomedical Engineering, State University of New York, Buffalo, NY 14260, USA; Medicine, State University of New York, Buffalo, NY 14260, USA; Clinical & Translational Research Center, Buffalo, NY 14260, USA

## Abstract

Besides acting as an immunological shield, the N-glycans of SARS-CoV-2 are also critical for viral life cycle. As the S2 subunit of spike is highly conserved across beta-coronaviruses, we determined the functional significance of the five ‘stem N-glycans’ located in S2 between N1098-N1194. Studies were performed with 31 Asn-to-Gln mutants, beta-coronavirus virus-like particles and single-cycle viral replicons. Deletions of stem N-glycans enhanced S1 shedding from trimeric spike, reduced ACE2 binding and abolished syncytia formation. When three or more N-glycans were deleted, spike expression on cell surface and incorporation into virions was both reduced. Viral entry function was progressively lost upon deleting the N1098 glycan in combination with additional glycosite modifications. In addition to SARS-CoV-2, deleting stem N-glycans in SARS-CoV and MERS-CoV spike also prevented viral entry into target cells. These data suggest multiple functional roles for the stem N-glycans, and evolutionarily conserved properties for these complex carbohydrates across human beta-coronaviruses.

**Author Summary:** Previous work shows that the N-linked glycans of SARS-CoV-2 are essential for viral life cycle. Few natural mutations have been observed in the S2-subunit of the viral spike glycoprotein in GISAID data, and mutations are absent in the five ‘stem N-glycans’ located between N1098-N1194. In the post-fusion spike structure these glycans lie equidistant, ~4 nm apart, suggesting functional significance. Upon testing the hypothesis that these glycans are critical for SARS-CoV-2 function, we noted multiple roles for the complex carbohydrates including regulation of S1-subunit shedding, spike expression on cells and virions, syncytial formation/cell-cell fusion and viral entry. Besides SARS-CoV-2, these glycans were also critical for other human beta-coronaviruses. Thus, these carbohydrates represent targets for the development of countermeasures against future outbreaks.

## Introduction

Beta-coronaviruses (β-CoVs) are enveloped, positive-sense single-stranded RNA viruses that cause respiratory diseases in humans (1, 2). Among these, OC43 and HKU1 cause mild infection, while zoonotic coronaviruses such as the 2002 SARS-CoV and 2012 MERS-CoV cause severe acute respiratory syndrome. More recently, the spread of the SARS-CoV-2 virus led to the COVID-19 pandemic. A common feature of these viruses is the spike glycoprotein, which is extensively decorated by a number of N- and O-linked glycans (3, 4). By binding various host cell receptors, especially angiotensin-converting enzyme 2 (ACE2) for SARS-CoV (5) and SARS-CoV-2 (6), and dipeptidyl peptidase-4 (DPP4, CD26) for MERS-CoV (7), spike mediates viral entry into host cells and also cell-cell transmission via syncytia formation (8, 9).

Whereas glycans on viral spike protein are traditionally thought to act as immunological shields that enable immune escape, other functions have also been attributed. Notably, truncation of both the SARS-CoV-2 spike glycoprotein N- and O-glycans using genetic methods reduced viral entry into human cells expressing ACE2 *ex vivo*, with N-glycans playing a more dominant role (10). This suggests functional roles for these complex carbohydrates in regulating viral entry functions. Consistent with this notion, treatment of these virus with peptide:N-glycanase (PNGaseF) (11, 12) and small molecule inhibitors of glycosylation (13, 14) also dramatically reduced viral entry into ACE2 expressing cells. Besides regulating viral entry, spike N-glycans are also thought to function by binding lectins such as C-type and Tweety family member 2 lectins on mononuclear blood cells to promote proinflammatory response (15). These glycans also engage host lectin receptors such as DC-SIGN (CD209), L-SIGN and Siglec-1 to promote viral attachment (16, 17). Additionally, the receptor-binding-domain (RBD) of spike is reported to contain a positively charged interface proximal to the ACE2 binding site that binds both heparan sulfate glycosaminoglycans (GAGs) (18, 19) and mono-sialylated glycolipids (20). These data suggest that glycans play essential roles in controlling viral function beyond functioning as an immunological shielding.

Studies focused on the effects of site-specific glycosylation using pseudotyped Vesicular Stomatitis virus (21) and lentivirus (22, 23) suggest that the modification of glycans at specific sites can reduce viral function though detailed mechanistic studies are not part of these investigations. Additionally, computational simulations propose that the glycans at N165, N234 and N343 within the spike N-terminus domain (NTD) and RBD may regulate the ‘up’ and ‘down’ conformation of RBD thus impacting receptor binding kinetics (24–26). Our prior studies also show that the N-glycans proximal to the S1/S2 polybasic cleavage site, in particular at N61 and N801, regulated spike incorporation into viral particles (12). Mutations at these sites impaired viral entry function. Moreover, bioinformatics analysis of N-glycosylation sites in GISAID (Global Initiative on Sharing All Influenza Data (27)) data suggests low mutation rates within spike as the virus evolves. This was particularly low among the N-glycans of the spike S2 subunit between N1098 and N1194 (12). These data suggest that N-glycans are essential and may have multiple effects on viral life cycle and entry function.

As the N-glycans in the stem region of the S2 subunit of spike have low mutation rates for SARS-CoV-2 and since they are conserved across human β-CoVs, this study determined their functional significance. In the case of SARS-CoV-2, these stem glycans lie at N1098, N1134, N1158, N1173 and N1194, and they lie equidistant (~4nm apart) in the spike postfusion structure (28, 29). Using a panel of Asn-to-Gln mutant spike in the context of virus-like particles (VLPs), single-cycle replicons and cell-based assays, we noted that these glycans are critical for the production of infectious virus. Deleting these glycans by introducing site-specific modification both reduced viral entry function and abolished cell-cell syncytia formation. In particular, we noted a prominent functional role for the N-linked glycan at N1098, which acted in synergy with other stem N-glycans especially N1173 and N1194. Besides impacting SARS-CoV-2 viral entry, ortholog N-glycans in the 2002 SARS-CoV and 2012 MERS-CoV also controlled viral entry into ACE2 and DPP4 expressing cells. Overall, our study provides mechanistic insight on the role of stem N-glycans in β-CoV life cycle and highlights the biological significance of these complex carbohydrates across human β-CoVs.

## Results

### Stem N-glycans are critical for cell surface expression of spike and ACE2-Fc functional binding

Natural mutations in SARS-CoV-2 spike are not common at sites of N-glycosylation (12). Among the 22 N-glycosylation sites on spike, loss of N-glycans due to modification of the N-X-S/T sequon have been noted at N17 in a number of variants of concern/ interest, and at N74 as part of the short-lived lambda strain (**Figure 1A**, **Supplemental Figure S1**) (30). Gain of glycan mutations were observed earlier in gamma at N20 and N188, but this was not sustained in subsequent lineages. More recently, gain of glycan mutations were reported due to H245N and K356T in the B.2.86 sub-lineage, and this is also observed in subsequent strains (31, 32). It is proposed that these novel gain-of-glycan mutations at N245 and N354 are a result of widespread use of COVID-19 vaccines which have strengthened the viral immunological shield (31, 32). Additionally, these mutations may also contribute to enhanced spike binding affinity (33). Mutations at the remaining N-glycosylation sites are relatively rare, particularly in the S2-subunit of spike.

**Figure 1.**
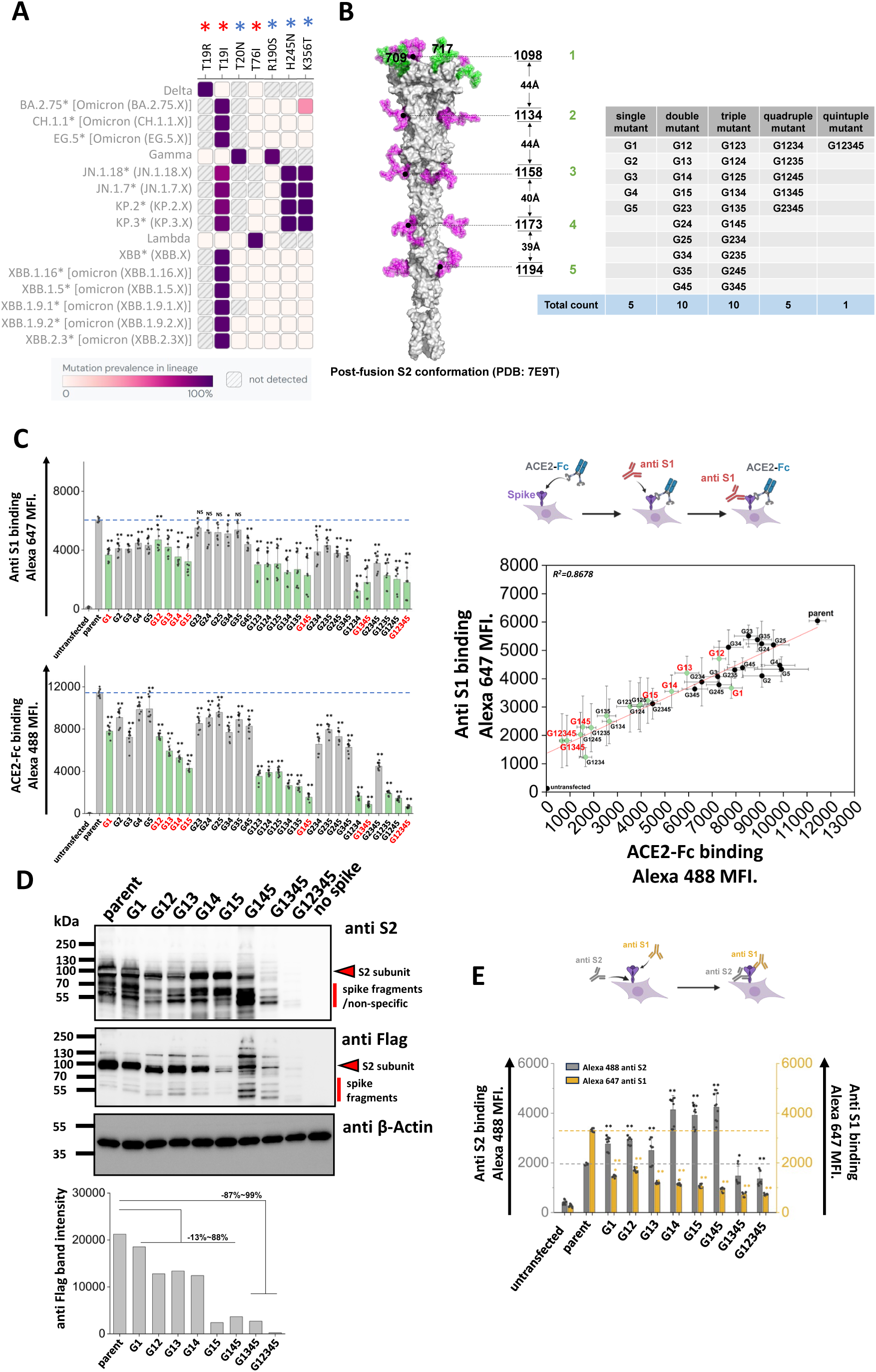
S2 stem N-glycans are critical for spike cell surface expression and ACE2-Fc functional binding. **A.** Heat map of mutation prevalence across SARS-CoV-2 world health organization variants of concern and interest. Red asterisk highlights glycosylation sites lost in individual viral strains and blue asterisk highlights the new glycosylation sites that have appeared. Data are rendered using dashboard at outbreak.info, using GISAID data. Only mutations with >75% prevalence in a single lineage are plotted. Each lineage is sequenced at least 1000 times. **B.** Post-fusion S2 conformation of spike protein (PDB: 7E9T) with five S2 stem N-glycans distributed along the axis. The distance between adjacent glycans is indicated. Asn (N) to Gln (Q) Spike N-glycan mutants at N1098Q, N1134Q, N1158Q, N1173Q and N1194Q are designated G1, G2, G3, G4 and G5, respectively. All possible N-to-Q mutant combinations were produced as shown in the table. **C.** 31 spike mutants and parent spike were transiently expressed in 293T cells. Anti-S1 mAb and ACE2-Fc fusion protein binding were simultaneously detected on cell surface. Mutations at N1098 (G1, shown using green bars) reduced spike function, particularly in synergy with additional N-to-Q mutations at other sites. ACE2-Fc binding was abolished in G1234, G1345 and G12345 mutants. The relationship between cell surface spike expression and ACE2-Fc binding was linear (R^2^ = 0.87). Selected G1-mutants, labeled red, were further analyzed in later studies. **D.** Western blots using anti-Flag and anti-S2 confirmed that stem N-glycan mutations reduce cell-surface expression, particularly upon implementing multiple edits. Anti β-Actin served as loading control. Note that some non-specific or Spike fragment bands appear when using cell lysates, but these are typically absent in virus blots. Densitometry was performed to quantify anti-Flag band intensity reduction. **E.** Anti-S1 and anti-S2 mAbs were applied in flow cytometry studies. The ratio of anti-S1 to anti-S2 mAb binding decreased in many cases suggesting enhanced S1 shedding upon implementing stem mutations. Data are Mean ± STD. * *P*<0.05, ** *P*<0.01, \*\*\**P*<0.001 with respect to parent.

As mutations in the stem N-glycans are infrequent, we tested the hypothesis that these carbohydrates may regulate spike function and be critical for SARS-CoV-2 viral life cycle. To test this, a panel of 31 spike mutants were created by implementing Asn-to-Gln (N-to-Q) mutation(s) combinatorially at positions N1098, N1134, N1158, N1173 and N1194 (**Figure 1B**) (28). These mutations were implemented on a base parent spike containing the dominant D614G mutation and C-terminus Flag-tag. Depending on the mutation site, these are abbreviated from G1 to G5. This panel includes five single, ten double, ten triple, five quadruple and one quintuple mutant that lacks all five stem N-glycans. In studies aimed at examining the effect of these site-specific glycan deletions, we noted that stem N-glycan mutations generally resulted in reduced S1-domain expression (measured using anti-S1 Ab) on the cell surface, and also reduced ACE2-Fc binding to cells (**Figure 1C**, **Supplemental Figure S2**). This was especially observed for either the single G1 (or N1098Q) mutation or other combinations that included G1 (green bars in **Figure 1C**). The reduction in anti-S1 binding correlated with ACE2-Fc binding. Loss of function was generally increased upon implementing more than one N-glycan deletion, with G12345 nearly abolishing both anti-S1 and ACE2-Fc binding. Overall, the stem N-glycans are critical for S1 cell surface expression and ACE2 receptor binding, with N1098 acting in synergy with other stem N-glycans, especially N1173 and N1194.

To investigate if the loss of S1 presentation was due to reduced protein expression, more detailed investigations were performed with selected spike mutants containing G1 (labeled red in **Figure 1C**). Cell lysates expressing these spike constructs were resolved using SDS-PAGE and probed with anti-S2, anti-Flag and anti-β-Actin antibodies in western blots (**Figure 1D**). Here, spike appears as a single ~95 kDa band as it was nearly completely cleaved within 293T cells at the furin site (10). The results showed that parent spike was efficiently expressed in cells. Spike mutants containing single G1, double and triple mutants (i.e. G145) exhibited 13~88 % decrease in intact S2 expression based on densitometry. Implementing quadruple (G1345) and quintuple (G12345) mutations resulted in more dramatic 87~99 % reduction in spike expression. Thus, the stem N-glycans may contribute to spike glycoprotein stability, particularly N1098 in synergy with other stem N-glycans.

As the decrease in ACE2-Fc binding in single and double site mutants (**Figure 1C**), was not accompanied by a proportional reduction in cellular spike expression based on western blots (**Figure 1D**), cytometry studies were undertaken to determine if mutations in stem glycans also result in enhanced S1 shedding (**Figure 1E**). To this end, the binding of anti-S1 and anti-S2 Abs to selected SARS-CoV-2 spike mutants was measured to compare S1 presentation with total spike expression on cells. In these studies, the ratio of anti-S1/anti-S2 binding decreased upon implementing single site-mutations, with this ratio being further reduced for the double and triple mutants. Whereas anti-S2 binding was equal to or higher than parent levels for the single, double and triple mutants, this was reduced for the quadruple and quintuple mutants, likely due to reduced spike stability as seen in the western blots in **Figure 1D**. The increased anti-S2 Ab binding in single, double and triple mutants may be a consequence of enhanced exposure of S2 subunit, following shedding of S1. Overall, the data suggest that single stem N-glycan mutations may promote the dissociation of S1 from spike, a process known as ‘shedding’. Implementing larger number of mutations may affect protein stability reducing spike protein expression on cell surface.

As glycans are essential for protein maturation, folding and intracellular translocation (12), we determined if protein instability induced by stem N-glycan mutations also promoted spike retention within cells. This was investigated using four-color imaging cytometry (**Supplemental Figure S3**). In the study design, FITC-anti-Flag antibody probed spike protein, Alexa 555-anti-Calnexin (CANX) antibody stained the endoplasmic reticulum (ER) (34), Alexa 647-anti-GM130 marked cis-Golgi (35) and Alexa 405-wheat germ agglutinin (WGA) was used to detect the cell membrane owing to its high affinity for diverse glycans (36). A gating strategy was implemented to select for single cells that were stained by all four markers (**Supplemental Figure S3A**). Representative images are displayed in **Supplemental Figure S3B** for the different spike mutants. Similarity analysis histograms quantified the co-localization coefficient between the different stains used in the study, including spike co-localization with different cellular organelle markers (**Supplemental Figure S3C**). Statistical analysis is presented in **Supplemental Table S1** for three different biological replicates. In all cases, a majority of spike signal co-localized with the ER and cis-Golgi markers. The measured signal with cell membrane was small, since WGA bound many components of the cell membrane glycocalyx, in addition to spike. Upon implementing stem N-glycan deletions, similarity score increased for spike co-localization with ER from 0.17 ± 0.02 for parent to 0.25 ± 0.05 for all stem N-glycan deletions. For cis-Golgi co-localization, these values increased from 0.94 ± 0.04 to 1.34 ± 0.18. The data suggest partial enhancement of spike retention in intracellular ER/Golgi compartments upon implementing glycan site-specific deletions.

Together, the data show that stem N-glycans regulate ACE2-Fc binding function with G1 mutations acting in synergy with other glycan deletions. In single, double and some triple mutants, the decreased function may be attributed to enhanced S1 shedding. In other triple, quadruple and quintuple mutants, protein misfold may occur resulting in reduced stability and expression on cell surface. The impact of glycan site-mutations on intracellular spike spatial distribution was small compared to their effect on shedding and cell surface expression.

### S2 stem N-glycans are critical for cell-cell syncytia formation

Besides direct viral entry, virus-induced cell-cell syncytia formation also contributes to transmission and disease pathogenesis (12, 37). This is a consequence of cell-cell fusion triggered by spike expressing infected cells fusing with neighboring ACE2 expressing cells, resulting in the formation of deletions reduce syncytia formation, we transiently expressed the spike mutants on 293T cells and mixed them with ACE2-expressing cells. Co-culture of cells resulted in syncytia formation, which was recorded using Incucyte live-cell imaging **(Figure 2A)**. Here, parent spike expressing cells consistently induced syncytia formation within 2 h post-mixing, with fusion area continuing to increase with time and cell rupture being observed when membranes were over-stretched **(Figure 2B, Supplemental Video S1-S5)**. The lack of just the N1098 glycan reduced syncytial area by >90 % 16 h post-mixing. Implementing more N-glycan mutations further reduced syncytia formation with complete abrogation in G12345. Overall, single-site mutations resulted in a more dramatic reduction in syncytia formation, compared to what would be anticipated based on partial reduction in S1 expression and ACE2-Fc binding (**Figure 1**). This suggests that the stem N-glycans may have additional effects in regulating cell-cell fusion.

**Figure 2.**
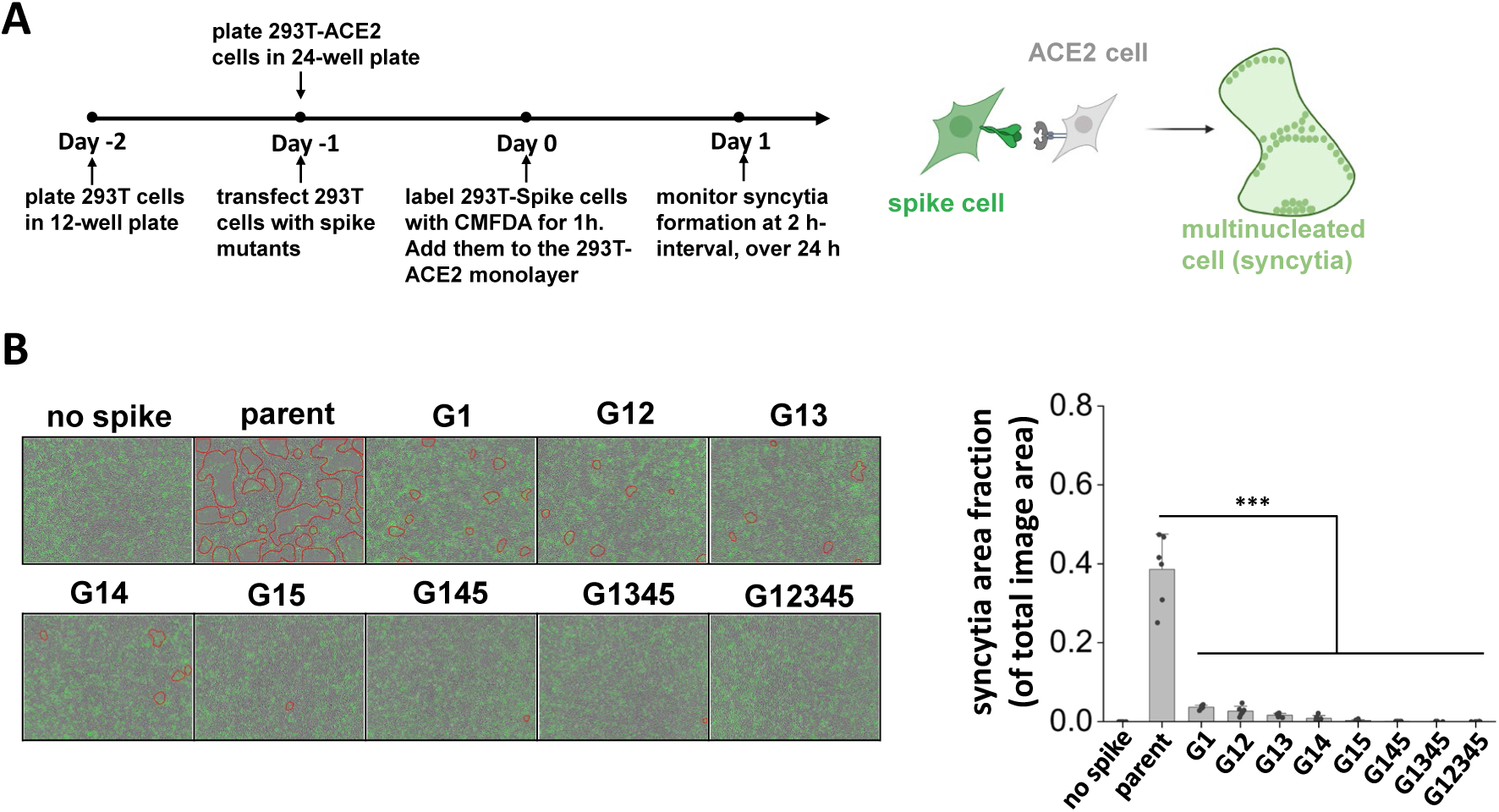
S2 stem N-glycans are critical for cell-cell syncytia formation. **A.** Schematic showing the Incucyte live-cell imaging workflow. 293T cells were transiently transfected to express selected spike mutants for one day, prior to cell labelling using 5-Chloromethylfluorescein diacetate, CMFDA. Spike expressing cells were then applied onto a monolayer of unstained 293T-ACE2 acceptor cells, and imaged every 2 h up to 24 h to measure syncytia formation. **B.** Representative images of syncytia formation at 16 h post-mixing are shown. Syncytia area was circled by red border. Syncytia area fraction = area occupied by syncytia/ total image area. All mutants demonstrated reduced syncytia formation, suggesting roles for S2 stem N-glycans in cell-cell viral transmission. Data are Mean ± STD. \*\*\**P*<0.001 with respect to parent.

### Stem N-glycans are critical for viral infection using SARS-CoV-2 virus-like particles (VLPs)

To investigate if the stem N-glycans affect SARS-CoV-2 viral infectivity, a ‘2-plasmid’ SARS-CoV-2 VLP system was developed. Here, the single plasmid (LVDP CMV-NME EF-1α-Luc-PS9) encoded for the SARS-CoV-2 nucleocapsid (N), membrane (M) and envelope (E) proteins along with firefly luciferase reporter gene complexed with viral RNA packaging signal ‘PS9’ **(Figure 3A)** (38). This vector was co-transfected along with spike expressing plasmid into 293T cells to produce ~100nm sized VLPs. VLPs with different spike mutants were produced in this manner and viral entry assayed using three target cell types, kidney 293T cells expressing ACE2 (293T-ACE2), lung epithelial A549-ACE2-TMPRSS2 cells which overexpress human ACE2 and TMPRRS2, and wild-type Calu-3 lung epithelial cells **(Figure 3B)**. Strikingly, we observed progressive reduction in viral entry upon implementing multiple glycan deletions with >95 % reduction being noted for G1345 and G12345 in all three target cell types. These data suggest that the stem N-glycans play pivotal roles in viral entry.

**Figure 3.**
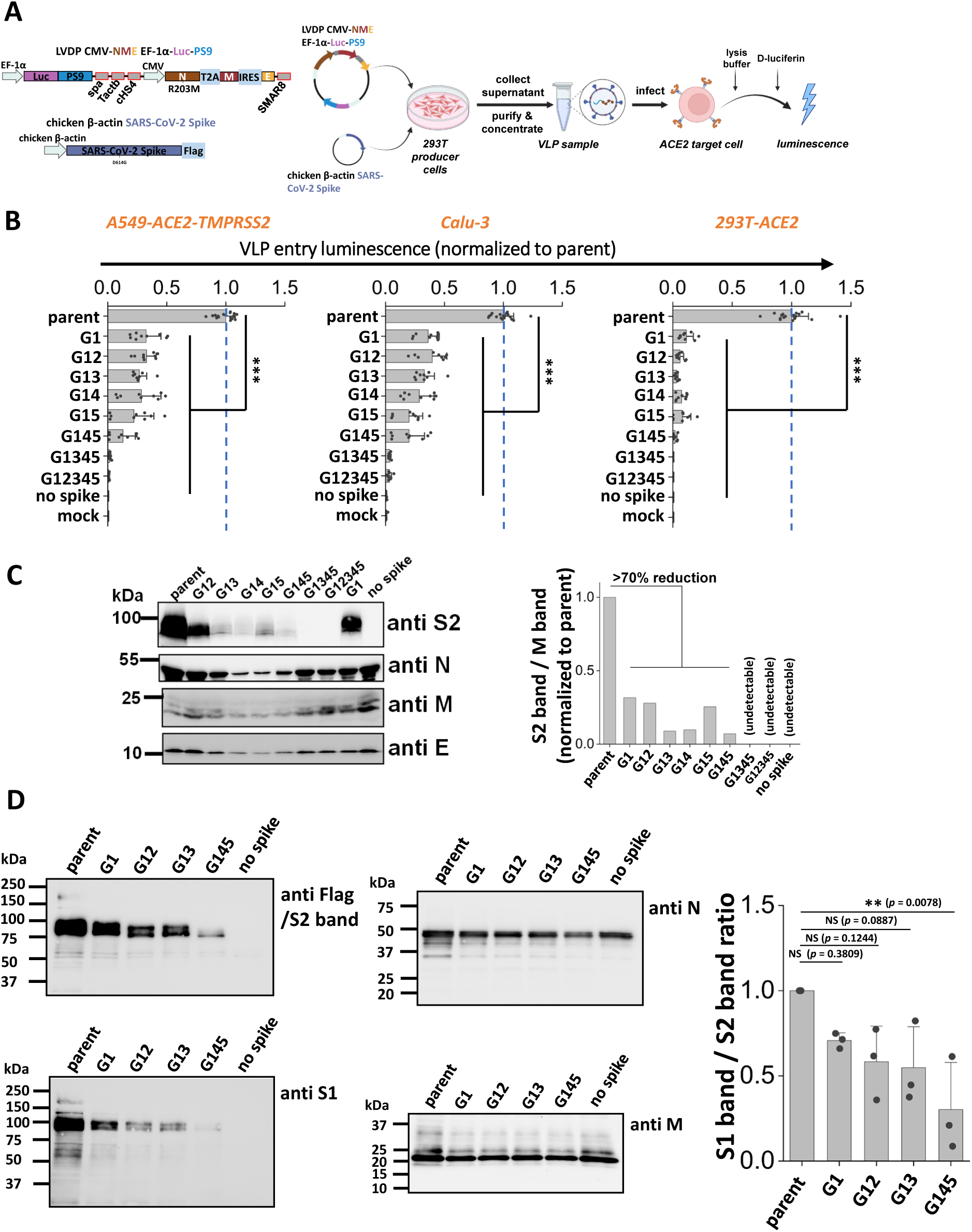
S2 stem N-glycans are critical for SARS-CoV-2 virus-like particle (VLP) infection. **A.** Workflow for producing SARS-CoV-2 VLPs using the 2-plasmid (‘2P’) system. VLPs are formed upon co-transfecting 293T cells with spike and LVDP CMV-NME EF-1α-Luc-PS9 plasmid. The latter construct contains two gene cassettes, one encoding for nucleocapsid (N), membrane (M) and envelope (E) proteins, and the second encoding luciferase with a cis-acting packaging signal ‘PS9’. 48 hours post-transfection, supernatant containing VLPs was harvested, clarified and concentrated. In functional assays, VLPs were added to target cells overnight, before measuring luciferase activity in cell lysate. **B.** Viral entry upon application of VLPs expressing different S2 mutants into three types of target cells, A549-ACE2-TMPRSS2, Calu-3, and 293T-ACE2. All G1 mutants displayed reduced viral entry, with greater reduction being observed upon incorporating more than one glycan mutation. **C.** Western blots of the VLPs. Densitometry was performed to normalize based on the M protein band. Spike incorporation into VLPs was reduced upon implementing S2 stem N-glycan mutations. **D.** Western blots of a sub-group of the selected spike mutant VLPs. Spike on VLPs were almost completely cleaved. S1 band/S2 band ratio was used for evaluating S1 shedding on VLP spike, with lower value indicating increased S1 shedding. A decreasing trend was observed as more stem N-glycans were deleted. VLPs Data are Mean ± STD. ** *P*<0.01, *** *P*<0.001 with respect to parent.

To determine how the stem N-glycans affect spike incorporation into VLPs, western blot analysis was performed for each of the VLPs containing mutant spike, using four antibodies that bind the spike S2 subunit (~95kDa), Nucleocapsid (~46kDa), Membrane (~25kDa) and Envelope (~10kDa) proteins. The results showed that the parent spike was efficiently incorporated into VLPs. The single and double mutants, G1 and G12, caused partial reduction in spike incorporation into VLPs. The remaining mutants displayed more dramatic reduction in spike incorporation **(Figure 3C)**. To quantitatively compare the band intensities, densitometry was performed by normalizing the anti-S2 band intensity based on the measured anti-M signal. While anti-M data are presented for such normalization, similar results were also noted upon using anti-N and anti-E as loading control. In such analysis, spike intensity varied as parent > G1 ~ G12 > other double and triple mutants. Spike was not incorporated in VLPs bearing G1345 and G12345, though clear bands were observed for the remaining structural proteins. To determine if S1 domain shedding from spike is augmented upon implementing stem N-glycan mutations, additional studies were performed with selected VLPs expressing G1, G12, G13 and G145 (**Figure 3D**). Upon comparing the intensity of anti-S1 band with respect to the anti-S2 band, we noted a progressive decrease in both bands upon implementing glycan mutations only the S1 band decreased more rapidly compared to the S2 band. This is particularly apparent upon performing densitometry analysis across multiple VLP batches (**Supplemental Figure S4**). In summary, stem N-glycan deletion reduced SARS-CoV-2 viral entry. This was partially due to reduced spike incorporation into VLPs and also due to enhanced shedding of the S1-subunit upon implementing these site-specific mutations.

### Stem N-glycans are critical for viral infection in studies using SARS-CoV-2 ΔS-virus-replicon-particles (ΔS-VRPs)

Although the SARS-CoV-2 VLPs carry the Luc-PS9 reporter that efficiently enables measurement of viral entry, it lacks a majority of the authentic SARS-CoV-2 viral genome. This may impact viral entry and host interaction features. To better mimic the authentic SARS-CoV-2 virion, single-cycle virus carrying spike glycan mutations were developed by adopting the SARS-CoV-2 ΔS-virus-replicon-particle (ΔS-VRP) system (39). This system contains the entire viral genome, only replacing the spike gene and a small 3’ portion of ORF1b with a Gaussia Dura-P2A-mNeonGreen reporter cassette **(Figure 4A)**. This construct is cloned into a bacterial artificial chromosome (bacmid) backbone. Transfection of host cells with the modified SARS-CoV-2 bacmid along with spike plasmid results in single-cycle non-replicative virions that can be used for viral entry investigations in BSL-2 setting.

**Figure 4.**
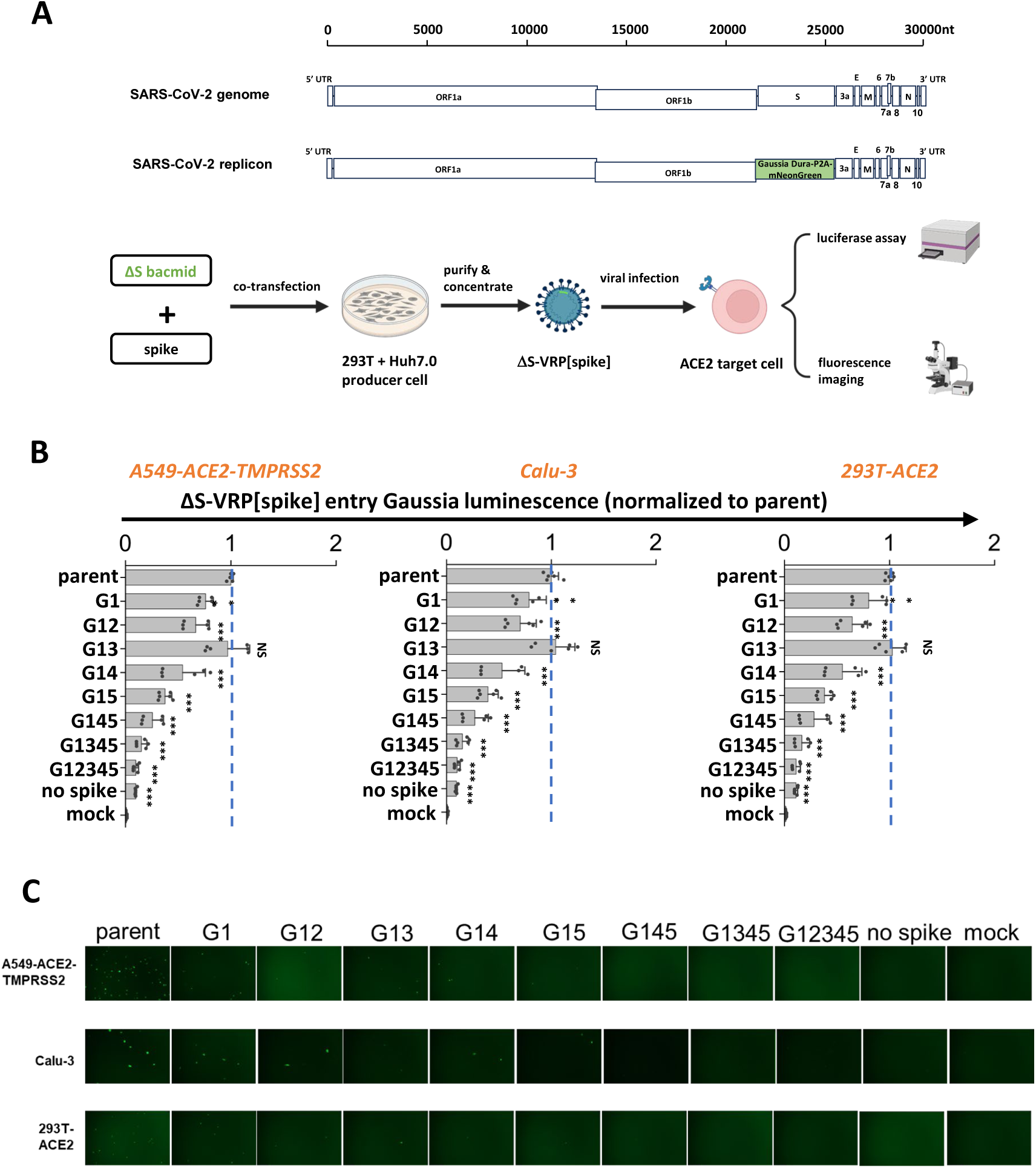
S2 stem N-glycans are critical for viral infection in assays using SARS-CoV-2 virus-replicon-particles (VRPs). **A.** Workflow for producing VRPs bearing SARS-CoV-2 spike glycoprotein. In the SARS-CoV-2 replicon, a Gaussia Dura-P2A-mNeonGreen reporter replaces spike gene along with a small 3’-portion of ORF1b. To produce the ΔS-VRP[spike], replicon bacmid and spike plasmid are co-transfected into pooled Huh7.0 and 293T cells. Supernatant harvested at 72h are concentrated to obtain VRPs. During functional studies, VRPs were added to target cells overnight, with viral entry being quantified both using fluorescence microscopy and Gaussia luciferase assays. **B-C.** ΔS-VRP[spike] expressing selected S2 mutants were added to three target cell types for viral infection assay.

Whereas the previous work demonstrated that the ΔS-VRPs could be trans-complemented with vesicular stomatitis virus G (VSV-G) glycoprotein, we extended this approach in the current manuscript by developing a protocol to enable SARS-CoV-2 spike incorporation into these single-cycle virions (details in Methods). Using this optimized system, parent and mutant spikes were successfully trans-complemented to make ΔS-VRP[spike]. The replicons with parent spike produced in this manner efficiently infected This was confirmed based on both a Gaussia luminescence assay (**Figure 4B**) and fluorescence microscopy (**Figure 4C**). While the spike G1 mutation partially reduced viral entry, implementing additional modifications particularly G14 and G15 further reduced viral infection. G145, G1345 and G12345 showed ~90 % reduction in viral infection, and almost no GFP positive cell in microscopy investigations. In negative controls, the measured signal was negligible in mock control and when ΔS-VRP were produced without spike. We note that ΔS-VRP[VSV-G] exhibited higher infectivity compared to ΔS-VRP[spike]. This is mainly due to the broad tropism of VSV-G which results in higher replicon titer production (**Supplemental Figure S5**). ΔS-VRP[spike] is produced at lower titer possibly due to syncytia formation and limited cell transmission in the producer cells that hampers virus generation. Regardless of this limitation, the data using replicons confirmed essential roles for stem N-glycans in regulating viral entry.

### Stem N-glycans are conserved, functional glycans in human beta-coronaviruses

Stem N-glycans are highly conserved across human β-CoVs, as noted upon sequence alignment of the S2 regions of common β-CoVs, including SARS-CoV-2, SARS-CoV, MERS-CoV, OC43 and HKU1 (**Figure 5A**). To determine if this evolutionary conservation has implications for viral function, studies were conducted with spike from 2002 SARS-CoV and 2012 MERS-CoV. Glycans in the stem region of these two proteins are shown in red or blue in **Figure 5A**. N-to-Q mutation was implemented at these sites to delete corresponding N-glycans. Thus, all five stem N-glycans of SARS-CoV were deleted to produce ‘SARS all5KO’. The seven stem N-glycans of MERS-CoV were divided into two groups, with the first 3 N-glycans being deleted in ‘MERS first3KO’, the remaining being deleted in ‘MERS last4KO’ and all 7 N-glycans deleted in ‘MERS all7KO’. Wild-type SARS (‘WT SARS’) and MERS (‘WT MERS’) were included as positive controls. VLPs were generated for each of these spike variants using the ‘2-plasmid’ system, and viral entry studies were performed using the relevant target cells **(Figure 5B)**. Here, the 293T-ACE2 cells were infected with SARS VLPs, while the 293T-DPP4 were infected with MERS VLPs, as the latter stably expresses DPP4/CD26. Viral entry results showed the absence of viral entry upon using SARS all5KO, >80 % reduction in viral entry using MERS first3KO, >90 % reduction for MERS last4KO and >95 % reduction for MERS all7KO. These observations establish a critical role for stem N-glycans in regulating viral infection across human β-CoVs.

**Figure 5.**
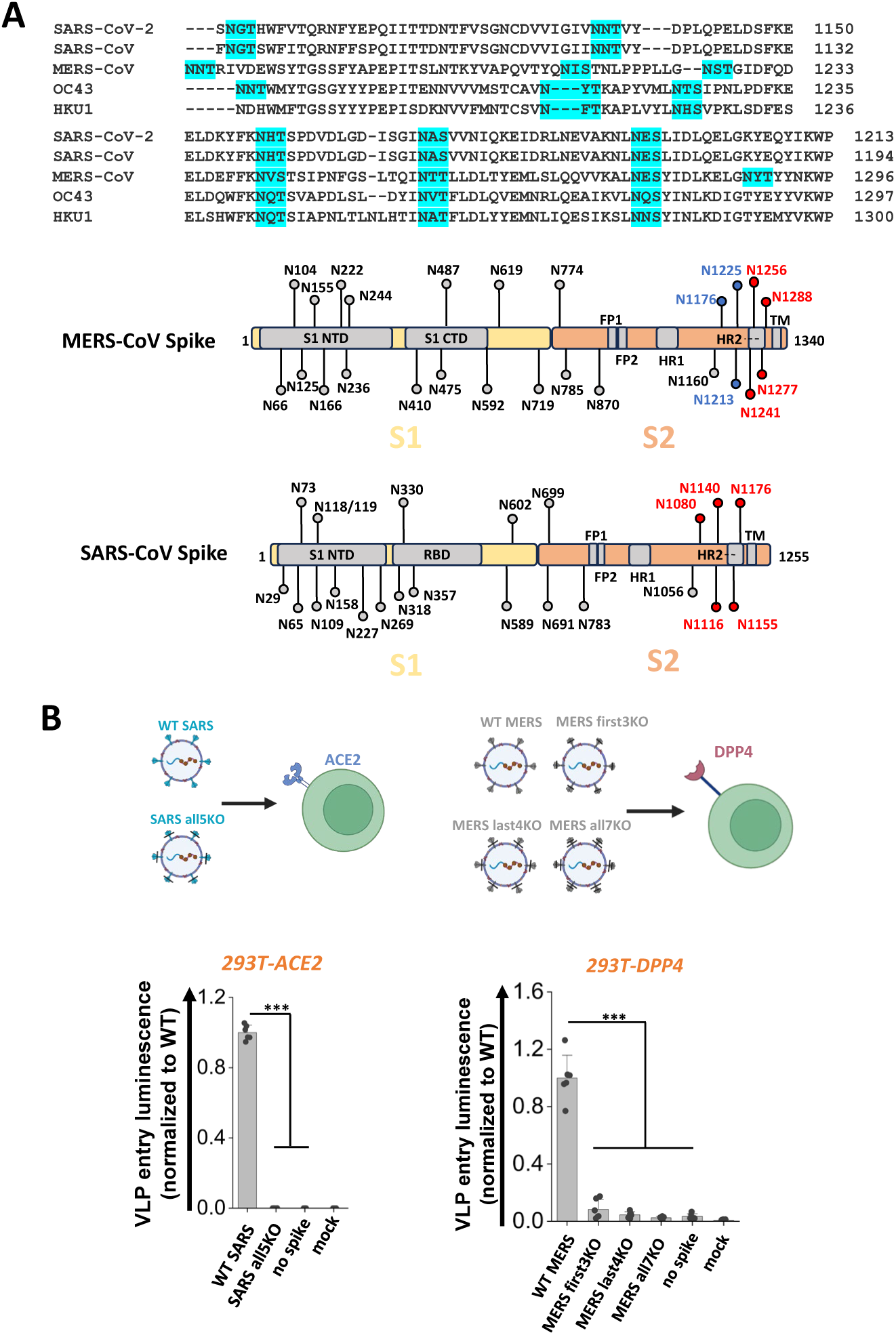
S2 stem N-glycans are conserved, functional glycans in all betacoronaviruses. **A.** Upper panel: sequence alignment of spike S2 stem region from five common human betacoronaviruses (hCoVs): SARS-CoV-2, SARS-CoV, MERS-CoV, OC43 and HKU1. The stem N-glycans are highlighted in cyan, and these are well conserved across betacoronaviruses. Lower panel: SARS-CoV and MERS-CoV were selected for further studies. The S2 stem N-glycans are colored in red or blue. **B.** All five S2 stem N-glycans of SARS-CoV were mutated N-to-Q to generate ‘SARS all5KO’. Three MERS mutants were also created upon implementing N-to-Q mutations at the first 3 glycosylation sites (‘first3KO’), last 4 sites (‘last4KO’) and all 7 glycosylation sites (‘all7KO’). VLPs made using SARS-CoV spike and MERS-CoV spike variants were used to infect 293T-ACE2 and 293T-DPP target cells, respectively. Viral entry was quantified using luminescence reporter. Absence of S2 stem N-glycans on all VLPs reduced viral infection. Data are Mean ± STD. \*\*\**P*<0.001

## Discussion

The continuous emergence of novel SARS-CoV-2 variants of interest (VOIs) and variants of concern (VOCs) underscores the importance of lasting virus surveillance and the need to expand our understanding of viral entry mechanisms (40). This is also necessary for determining pan-coronavirus inhibition strategies, in preparation for future infections and disease. The S2 stem region of coronavirus spike stands out as an attractive target for such therapeutics due to its striking evolutionary conservation (41, 42). Consistent with this, our previous bioinformatics analysis suggests very low number of glycan mutations in this region (12). Thus, this conserved region along with the stem N-glycans would be an attractive target for limiting β-CoVs related diseases. To investigate this, we created 31 SARS-CoV-2 spike mutants that lack various combinations of the conserved stem N-glycans. Our studies reveal multiple roles for these complex carbohydrates.

In one aspect, we observed that mutations in stem N-glycans may augment S1 subunit shedding, and this directly correlated with the ability of spike to bind ACE2. S1 shedding was also observed in the mutant VLPs and this contributed to reduced viral entry. Related to this, we previously reported that truncation of N-glycan biosynthesis at the high-mannose stage may increase spike proteolysis and shedding of S1 subunit (10), though the precise contributors were unclear. This is functionally important as others have demonstrated a correlation between the degree of S1 presentation and viral infectivity (43). Thus, spike cleavage at the furin site while promoting S2’ proteolysis and viral entry, also simultaneously limits viral entry by reducing virus binding to ACE2. In this current study, also, we noted a strong correlation in that glycan mutations that enhanced S1 shedding also proportionally reduced both VLP and VRP entry into a variety of host cells. In particular, the glycan at N1098 acted in synergy with other stem glycans, especially N1173 and N1194, to regulate both ACE2 binding and viral entry. Computational studies in literature suggest mechanisms supporting our wet-lab observations. Serapian *et al.* showed that carbohydrates including the stem N-glycans may exhibit strong energetic coupling to other regions of the protein, enhancing intramolecular interaction networks that stabilize spike (44). Teng *et al.* show that single point mutations at selected glycosites including N1098 may lead to spike instability (45). Together these stem N-glycans may contribute to the spike pre-fusion structure, potentially impeding S1 shedding and maintaining ACE2 binding function.

While mutations at single sites promoted shedding, multiple stem glycan site deletions resulted in reduced spike translocation onto both host cell surface and incorporation into virions. Several processes could contribute to these observations, including spike misfolding due to lack of interaction with intracellular chaperones like calnexin and calreticulin (12). This could then lead to either premature protein misfolding, intracellular retention or lack of spike trimerization (46). In this regard, indeed, our previous work demonstrated that spike glycans bind calnexin within cells and this is necessary for the production of functional virions (12). Our newer imaging cytometry studies add to this knowledge, suggesting only partial effects of stem N-glycans in regulating spike intracellular ER and Golgi retention. Related to this, Huang *et al*. report that the N1194Q mutation of spike partially disrupts spike trimerization resulting in expression of spike monomer protein in *in vitro* assays (22). Overall, implementing multiple stem glycan deletions resulted in defective spike expression, preventing spike incorporation into virions and reduced viral entry function.

Strikingly, whereas multiple glycan mutations were necessary to abrogate spike-ACE2 binding and viral infectivity, a single N1098Q mutation reduced syncytial formation by >90 %. This suggests additional roles for the stem glycans in mediating cell-cell fusion other than the pathways stated above. In agreement, Dodero-Rojas *et al.* (47) showed that the N-glycans in the stem region form a ‘glycan cage’ once the S1 subunit is shed from spike in the S2’ cleaved state. This cage structure impedes the movement of the stalk region of spike, leading to improved kinetic stability of spike. This improved stability promotes fusion peptide integration with target cell membrane, resulting in increased cell-cell fusion event occurrence (47). Disruption of glycan structures disrupts cage formation resulting in kinetic instability of the S2 stem fusion peptide, which then hinders the fusion process. Our wet-lab studies support these observations and suggest that N1098 may be a key carbohydrate in the glycan cage that is essential for structural arrangements that accompany cell-cell fusion.

The stem N-glycans are highly conserved in SARS-CoV-2 variants, even under natural selection. While most of the original 22 spike N-glycans have remained, some losses/gains have been reported in the S1 subunit glycans but not in the S2 glycans. In this regard, Alpha and Beta maintained the native glycans of the original virus, while Gamma gained two N-glycans at N20 and N188 (30). Delta and Omicron lost a single N17 glycosylation site with Lambda discarding N74. More recently, two new N-glycans (N245, N354) have been acquired after the B.2.86 sub-lineage in JN and KP strains, and this is thought to contribute to both augmented viral immunological shield and increased fitness (31, 32). In addition to glycosylation sites, even the glycoforms in the S2 subunit are largely conserved, with the stem N-glycans remaining mostly as complex N-linked carbohydrates through the course of evolution (3, 48, 49). On the other hand, selected N-glycans in the S1 domain that regulate spike-ACE2 interactions, specifically N165, N343 and N616, are reported to now appear in less processed high-mannose form in the newer virus strains (49). Changes in these key residues to mannose-rich form may contribute to reported enhanced susceptibility of Omicron to the potent Mannosidase-I inhibitor-drug kifunensine, compared to ancestral SARS-CoV-2 (50, 51). Thus, we speculate that in addition to glycosylation site, conservation of glycoforms in S2 may also be critical for virus function.

In addition to SARS-CoV-2, the stem N-glycans were also critical for SARS-CoV and MERS-CoV function, suggesting evolutionary conserved roles for these carbohydrates. While our studies focused on SARS-CoV-2 VLPs, future work may determine if the same is observed upon creating strain-specific viral replicons, and in studies that examine the impact of each of the glycosylation sites individually and synergistically on β-CoV function. In addition to human coronavirus, our observations may also be more broadly applicable to sarbecoviruses in animal reservoirs as well. In this regard, Allen *et al.* (52) reported that 15 of the 22 N-glycans of SARS-CoV-2 are shared by 78 different sarbecoviruses including all five stem N-glycans. Additionally, the N-glycans in the stem region were complex-type in all twelve sarbecovirus strains analyzed using mass spectrometry. Greater structural variation is noted in S1 subunit glycans, particularly those in the N-terminal domain (NTD). These findings are highly consistent with our proposition related to the evolutionary conserved functional roles for these N-glycans in coronavirus life cycle.

In summary, our data suggest that both the sites of N-glycosylation in the S2 stem region and glycoforms present there are highly conserved among β-CoVs. These carbohydrates are functionally critical for SARS-CoV-2, SARS-CoV and MERS-CoV. By acting in synergy, these glycans regulate multiple biological pathways. Such evolutionary conservation could serve as a motivation to develop pan-coronavirus countermeasures directed against these sites.

## Materials and Methods

### Materials

Recombinant human angiotensin-converting enzyme 2-Fc (ACE2-Fc) fusion protein was produced as previously described (10). Alexa 647-conjugated mouse anti-SARS-CoV-2 Spike S1 subunit mAb (IgG_1_, Cat#: FAB105403R), mouse anti-SARS-CoV-2 spike S2 subunit mAb (IgG2a, Cat#: MAB10557) and mouse anti-SARS-CoV-2 nucleocapsid protein mAb (IgG_2b_, Cat#: MAB10474) were from R&D Systems (Minneapolis, MN). Mouse anti-SARS-CoV-2 membrane protein mAb E5A8A (IgG1, Cat#: 15333), rabbit anti-SARS-CoV-2 envelope protein polyclonal antibody (pAb) (Cat#: 74698), rabbit anti-β-Actin mAb (IgG, Cat#: 8457), Alexa 647-conjugated rabbit anti-GM130 mAb (IgG, Cat#: 59890), Alexa 555-conjugated rabbit anti-Calnexin (CANX) mAb (IgG, Cat#: 23198), HRP-conjugated horse anti-mouse pAb (IgG, Cat#: 7076) and HRP-conjugated goat anti-rabbit pAb (IgG, Cat#: 7074), were from Cell Signaling (Danvers, MA). HRP-conjugated rat anti-Flag mAb (IgG_2a_, Cat#: 637311) was from BioLegend (San Diego, CA). FITC-conjugated mouse anti-Flag mAb (IgG1, Cat#: F4049) was from Millipore Sigma (Burlington, MA). Alexa 488-conjugated goat anti-human pAb (IgG, Cat#: 109-545-190) was from Jackson ImmunoResearch (West Grove, PA). In some cases, antibodies or lectins were conjugated with AZDye NHS ester (VectorLabs) by incubating 0.5-1 mg/mL protein in PBS (1 mM KH_2_PO_4_, 155 mM NaCl, 3 mM Na_2_HPO_4_) with 20 molar-fold excess AZDye NHS ester dye dissolved in DMSO for 1h at room temperature (RT). Reaction volume varied from 50-100 μl typically. Following reaction, 1/10^th^ volume 1M Tris was used to quench the reaction and the unreacted dye was removed using a 7 kDa cutoff Zeba desalting spin column that was equilibrated with PBS (ThermoFisher). All other biochemicals were from ThermoFisher (Waltham, MA), Sigma Chemical company (St. Louis, MO), or Vector laboratories (Newark, CA) unless otherwise mentioned.

### Molecular biology

The parent spike (full-length SARS-CoV-2 spike protein with C-terminal Flag-tag containing D614G mutation) was from previous work (12). A panel of 31 spike mutants lacking various combinations of N-glycans were created on this background by implementing site-specific Asn-to-Gln (N-to-Q) mutations. The LVDP CMV-NME EF-1α-Luc-PS9 plasmid used for SARS-CoV-2 virus-like particle (VLP) production will be described elsewhere (Yang *et al.*, unpublished data). The bacmid encoding the SARS-CoV-2 replicon was described previously (*39*). The pCDNA3.3 MERSD12 spike plasmid encoding the MERS WT spike protein with a 12-amino acid deletion at the C-terminal tail was a gift from Dr. David Nemazee (RRID: Addgene<u>_</u>170448). For simplicity, this protein is referred to as ‘MERS’ in this manuscript. The pcDNA3.1 SARS spike plasmid encoding the 2002 SARS spike protein was kindly provided by Dr. Fang Li (RRID: Addgene_145031). The pLEX307-DPP4-puro plasmid encoding DPP4/CD26 was a gift from Drs. Alejandro Chavez & Sho Iketani (RRID: Addgene_158451).

### Cell culture

Human embryonic kidney 293T Lenti-X cells (‘293T’) (Cat#: 632180) were purchased from Clontech/Takara Bio (Mountain View, CA). Stable 293T-human ACE2 (293T-ACE2) cells were kindly provided by Dr. Michael Farzan (Scripps Research, Jupiter, FL). 293T-DPP4 cells were generated by transducing lentivirus packaged with Dipeptidyl peptidase-4 (DPP4) gene into 293T cells, and subsequently culturing isogenic clones. Human adenocarcinoma alveolar basal epithelial A549 cells overexpressing ACE2 and TMPRSS2 (‘A549-ACE2-TMPRSS2’) (Cat#: a549-hace2tpsa) was purchased from Invivogen (San Diego, CA). Human airway epithelial Calu-3 (Cat#: HTB-55) was from ATCC (Manassas, VA). Hepatocyte-derived carcinoma Huh7.0 cells were available from our prior work (*39*).

### Transfection

293T cells were transfected via a calcium phosphate method described previously (53) or Lipofectamine 2000 reagent following manufacturer’s instructions. These transfections were used for transient expression of a panel of spike mutants in 293T cells. In brief, 1 million 293T cells were plated in 6-well plates one day prior to transfection. The next day, when cell density reached ~70 % confluence, 2 μg DNA was used to transfect each well in a 6-well plate. 6-8 h post-transfection, media was switched to fresh Opti-MEM (ThermoFisher).

### Flow cytometry

Cells transfected with spike were trypsinized from 6-well plates and resuspended in HEPES buffer (110 mM NaCl, 10 mM KCl, 2 mM MgCl_2_, 10 mM Glucose, 30 mM HEPES, pH = 7.2-7.3) at 10^7^/mL. 20 μl cells were then added into 1.5 mL eppendorf tubes along with fluorescent antibodies indicated in relevant figure legends at manufacturer’s recommended concentration. In some runs, ACE2-Fc fusion protein was added as described previously (10). The cells were then incubated for 15-20 min. on ice with periodic flicking, washed and resuspended at 2×10^6^/mL in HEPES, and analyzed using a BD Fortessa X-20 flow cytometer (San Diego, CA). Mean fluorescence intensity (MFI) was recorded.

### Western blot

VLP samples or cell lysate were prepared in SDS-DTT blue loading buffer (Cell Signaling) following manufacturer’s instructions and denatured at 98 ⁰C for 5-10 min. 10 μl VLP sample for anti-M and anti-E or 2 μl VLP sample for anti-S2 and anti-N, or 1 μl cell lysate sample for anti-Flag, anti-S2 and anti-β-Actin were resolved using a 12 % Tris-glycine gel. Following transfer onto a nitrocellulose membrane using a Trans-Blot Turbo Transfer System (Biorad, Hercules, CA), membranes were blocked in TBST (100 mM sodium chloride, 20 mM Tris-HCl, 0.1 % Tween-20) containing 5 % non-fat milk for 1-2 h at RT. The membranes were then incubated with primary antibody at recommended concentrations in TBST containing 2 % non-fat milk at 4 °C overnight. The next day, the membranes were washed with TBST four times with each wash lasting 5 min. at RT. The membranes were then, as necessary, treated with HRP conjugated secondary antibody for 1 h at RT at manufacturer recommended concentrations. Subsequently, the membranes were washed again using TBST solution four times with each wash lasting 5 min. at RT. In the final step, signal was developed using SuperSignal chemiluminescence substrate (ThermoFisher) and imaged using a ChemiDoc Imaging System (Biorad).

### Imaging cytometry

293T cells transfected with spike were resuspended in HEPES buffer at 10^7^ /mL. 1 μg/μl Alexa 405-conjugated wheat germ agglutinin (WGA) lectin was added into 400 μl spike expressing cells for 20 min. on ice. The cells were then fixed using 1.5 % paraformaldehyde for 1 h at RT, washed using 200 μl HEPES buffer and permeabilized using 200 μl ice cold pure methanol for 5-10 min. at 4 °C. Following permeabilization, the cells were washed using HEPES buffer and then incubated with Alexa 647-anti-GM130, Alexa 555-anti-Calnexin (CANX) and/or FITC-anti-Flag antibodies (to label spike) for 20 min. on ice at manufacturer recommended concentrations. Following incubation, cells were again washed with HEPES buffer and analyzed using a Cytek Amnis MKII Imaging cytometer (Fremont, CA).

The IDEAS Analysis Application v6.0 software was used for similarity score quantification. First, gates were set on the singlet cells to obtain cell populations that positively stained for all four fluorescent markers. Next, a threshold was set using the ‘mask’ function of the software to identify pixels in the image cytometry data that correspond to individual fluorophores. A single threshold setting was applied to all images collected in a single run. Representative images following thresholding for each of the spike mutants is presented in Supplemental Material. Finally, the built-in function ‘Similarity’ was utilized to determine the co-localization between different fluorescent regions by comparing the signal intensity in the different masks.

### Syncytia formation

293T cells were transfected with spike on day −1 (‘minus one’). 0.2-0.4 million 293T-ACE2 cells were also plated in 24-well plates on day −1. The next day, the 293T spike donor cells were labelled green using 5-Chloromethylfluorescein diacetate CellTracker CMFDA green dye (ThermoFisher) for 1 h in incubator following reagent’s manual. The spike donor cells were then washed using HEPES buffer once, trypsinized and resuspended in DMEM. 0.2-0.4 × 10^6^ of spike donor cells were then applied onto the monolayer of 293T-ACE2 acceptor cells. Immediately, the plate was placed in an Incucyte S3 Live-Cell Analysis System (Sarorius, Germany) and imaged at 2 h intervals for up to 24 h. Data were processed using ImageJ, with syncytia area being manually marked and counted. Syncytia area fraction = area occupied by syncytia/ total image area.

### SARS-CoV-2 virus-like particle (VLP) production

VLPs were produced using a 2-plasmid (“2P”) system where one plasmid expressed spike and the second plasmid (’ LVDP CMV-NME EF-1α-Luc-PS9’) expressed all other structural components along with luciferase reporter. Here, 15-20 × 10^6^ 293T cells were plated in 150 mm tissue-culture treated petri dishes. The next day, cells at ~70 % confluence were co-transfected with 50 μg LVDP CMV-NME EF-1α-Luc-PS9 and 1 μg spike plasmid using the calcium phosphate method. 6-8 h post-transfection, cell culture medium was switched to 20 mL fresh Opti-MEM. The cells were further incubated for 48 h to allow VLP production. Subsequently, the supernatant was collected, centrifuged at 4000 g for 5 min and filtered through a 0.45 μm polyethersulfone (PES) membrane to remove cell debris. The filtrate was then added to a polycarbonate centrifuge tube (Beckman Coulter, Indianapolis, IN). 20 % (g/mL) sucrose solution was loaded into the bottom of the tube using a 4’’ long stainless-steel needle, with the sucrose cushion volume being equal to 10 % of supernatant volume. This sample was then ultracentrifuged at 150,000 g for 2.5 h at 4 °C using a Type 70 Ti or Type 45 Ti rotor in an Optima XE Ultracentrifuge (Beckman Coulter). The translucent VLP pellet formed by this process was resuspended in 200 μl PBS buffer, placed in a 1.5 mL Eppendorf tube, vortexed thoroughly and spun down in a bench-top centrifuge at 13,000 g for 2 min. to remove any residual debris. The clear supernatant was then transferred into a new 1.5 mL Eppendorf tube, which was either directly used for viral entry assay or stored at −80 °C for future studies.

### SARS-CoV-2 virus-replicon-particle (VRP) production

The SARS-CoV-2 replicon bacmid was from our previous study (*39*). To produce the VRPs, 12 µg replicon bacmid DNA and 4 µg spike or VSV-G plasmid were mixed well in 500 µl serum reduced OPTI-MEM (Invitrogen). Simultaneously 48 µl lipofectamine 2000 was diluted in 500 µl serum reduced OPTI-MEM. The two samples were then mixed and incubated at room temperature for 15-20 min. During plasmid incubation, 4×10^6^ Huh7.0 cells and 4×10^6^ 293T cells were mixed and added into 100 mm petri dishes with 15 ml DMEM containing 10 % FBS. The DNA/lipofectamine 2000 mix was then added dropwise to cells with gentle rocking. 6-8 h post transfection, media was switched to 15 ml DMEM containing 2 % FBS and cells were further incubated for 72 h for VRP production. At the end point, supernatant containing VRPs was harvested, centrifuged at 4000 g for 5 min., filtered using 0.45 µm PES filters to remove cell debris, and then buffer exchanged to HEPES buffer using a 100 kDa PES protein concentrator (ThermoFisher). In addition to VRP preparation, this step also simultaneously depleted any residual luciferase signal from producer cells. The final VRP product was 30-fold concentrated in ~500 μl volume, and available for either immediate use in viral entry assays or stored at −80 °C for later use. The replicon particles resulting from the above steps are called ΔS-VRP[spike] if they bear SARS-CoV-2 spike glycoprotein or ΔS-VRP[VSV-G] if they are decorated by VSV-G.

### Viral entry luminescence assay

Target cells were trypsinized and resuspended at 10^7^ /ml. In a typical viral entry assay, 50 μl VLPs or ΔS-VRPs was mixed with 80,000 cells (8 μl of stock) along with 8 μg/mL polybrene in a 1.5 mL Eppendorf tube. This was kept at RT for 25 min. with periodic flicking. The cells were then added into 96-well plates and incubated overnight to allow the expression of reporter protein(s).

In the case of SARS-CoV-2 VLPs, firefly luciferase signal was measured based on previous work (12). In brief, the cells were washed with 200 μl PBS and lysed using 50 μl/well cell lysis buffer (Gold Biotechnology) at RT for 20 min. In the meantime, fresh 2X TMCA buffer was made by mixing 20 μl MgCl_2_ (from 500 mM MgCl_2_ 100X stock), 20 μl Coenzyme A (from 25 mM 100X stock), 20 μl ATP (from 15 mM 100X stock), 500 μl Tris-HCl, pH = 7.8 (from 400 mM 4X stock) and additional 440 μl cell culture water to bring up the volume to 1 mL. 50 μl cell lysate was then added into 96-well white plates with round bottom followed by addition of 50 μl 2X TMCA. 1 μl D-Luciferin (from 15 mg/ml D-Luciferin 100X stock) was then added and luminescence was immediately read using a BioTek Synergy4 plate reader (Santa Clara, CA).

In the case of SARS-CoV-2 replicons (ΔS-VRPs), secreted Gaussia-Dura luciferase was measured using the Gaussia Luciferase glow assay kit (ThermoFisher). Briefly, this involved addition of 50µl culture supernatant into white 96-well round bottom plates, along with 50µl working solution from the kit. Samples were incubated for 10 min. at RT to stabilize the luminescence signal before luminescence measurement.

### Biohazard

All protocols described above were conducted in BSL-2 facility, as approved by the University at Buffalo Biosafety Committee.

### Statistics

All data are presented as mean ± standard deviation for multiple biological replicates. Multiple comparisons were performed using ANOVA followed by the Student-Newman-Keuls post-test. **\****P*<0.05, **\*\****P*<0.01 and **\*\*\****P*<0.001 was considered to be statistically significant. Number of repeats are presented using discrete points in individual plots. Typically, three identical samples were measured together on the same day to account for technical variability, and different virus/cell batches prepared on different days accounted for biological reproducibility.

## Acknowledgments

We are grateful to the Cell, Gene and Tissue Engineering Center, University at Buffalo, for generous access to the Incucyte incubator-microscope system. This work was supported by a University at Buffalo Blue Sky award (S.N.), and NIH grants UL1TR001412 & HL103411 (S.N.). Imaging cytometry studies performed at the Roswell Park Comprehensive Cancer Center (RPCCC) Flow and Image Cytometry Shared Resources (FICSR) were partially supported by NCI grants P30CA01656 and NCI R50 R50CA211108.

## Data sharing plan

All data are presented in main figures and Supplemental data. Plasmid reagents are deposited at Addgene. Other reagents will be provided by the corresponding author upon request.

## Author contributions

Conceptualization: Q.Y., B.M. and S.N. Methodology: Q.Y., A.K., B.M., S.N. Investigation: Q.Y., S.N. Visualization: Q.Y. and S.N. Writing (original draft): Q.Y. Writing (review and editing): all authors.

